# Large RNP granules in *C. elegans* oocytes have distinct phases of RNA binding proteins

**DOI:** 10.1101/2022.05.25.493450

**Authors:** Mohamed T. Elaswad, Brooklynne M. Watkins, Katherine G. Sharp, Chloe Munderloh, Jennifer A. Schisa

**Affiliations:** Central Michigan University, Biochemistry, Cell and Molecular Biology Program; Central Michigan University, Department of Biology; Department of Biological Sciences, University of Pittsburgh; Department of Molecular and Human Genetics, Baylor College of Medicine

## Abstract

The germ line provides an excellent *in vivo* system to study the regulation and function of RNP granules. Germ granules are conserved germ line-specific RNP granules that are positioned in the *C. elegans* adult gonad to function in RNA maintenance, regulation, and surveillance. In *C. elegans*, when oogenesis undergoes an extended meiotic arrest, germ granule proteins and other RNA binding proteins assemble into much larger RNP granules whose hypothesized function is to regulate RNA metabolism and maintain oocyte quality. To gain insight into the function of oocyte RNP granules, in this report we characterize distinct phases for four protein components of RNP granules in arrested oocytes. We find the RNA binding protein PGL-1 is dynamic and has liquid-like properties, while the intrinsically disordered protein MEG-3 has gel-like properties, similar to the properties of the two proteins in small germ granules of embryos. We find that MEX-3 exhibits several gel-like properties but is more dynamic than MEG-3, while CGH-1 is dynamic but does not consistently exhibit liquid-like characteristics and may be an intermediate phase within RNP granules. These distinct phases of RNA binding proteins correspond to, and may underlie, differential responses to stress. Interestingly, in oocyte RNP granules MEG-3 is not required for the condensation of PGL-1 or other RNA binding proteins, which differs from the role of MEG-3 in small, embryonic germ granules. Lastly, we show the PUF-5 translational repressor appears to promote MEX-3 and MEG-3 condensation into large RNP granules; however, this role may be associated with regulation of oogenesis.

## INTRODUCTION

Germ granules are largely conserved in germ lines across metazoa. They are essential for germ cell differentiation and fertility, and germ granules appear to be the site where small RNA pathways monitor germ line transcripts (Kawasaki et al. 2004; Spike et al. 2008; Billi et al. 2012; Pek et al. 2012; Updike et al. 2014; Trcek and Lehmann 2019; Sundby et al. 2021). Due to their more complex *in vivo* environment, germ granules differ from *in vitro* condensates that are simple, single-phase liquid droplets. *Drosophila* germ granules (polar granules) contain multiple homotypic clusters of RNAs within protein-rich condensates (Little et al. 2015; Trcek et al. 2015). Similarly, in zebrafish and *Xenopus* an amyloid-like scaffold organizes RNAs into specific subregions within germ granules (Boke et al. 2016; Fuentes et al. 2018; Roovers et al. 2018).

In the early *C. elegans* embryo, germ granules (P granules) also have multiple phases. The intrinsically disordered protein MEG-3 is concentrated at the periphery of granules, while the RGG-domain protein PGL-3 is detected in the central core of granules (Wang et al. 2014; Putnam et al. 2019). Moreover, the MEG-3 phase has gel-like properties in terms of its dynamics and stability, whereas PGL-1 and PGL-3 are liquid-like and exhibit a sensitivity to temperature change (Brangwynne et al. 2009; Wang et al. 2014; Saha et al. 2016; Putnam et al. 2019). MEG-3 and its paralog MEG-4 are required to promote the condensation of PGL-3 (Putnam et al. 2019). Thus, it is well established that embryonic germ granules have multiple phases and a complex structure under physiological conditions. In young *C. elegans* hermaphrodites meiotic maturation and gamete production occur in an efficient, assembly line-like fashion; however, when sperm become depleted within a few days of adulthood, an extended meiotic arrest occurs where oocytes stack up in the adult gonad (McCarter et al. 1999). Several reversible, cellular changes in meiotically arrested oocytes have been documented, including reorganization of the microtubule cytoskeleton, remodeling of the endoplasmic reticulum, and the condensation of mRNA and RNA binding proteins into large RNP granules (Harris et al. 2006; Langerak et al. 2019; Schisa et al. 2001). The large RNP granules have been hypothesized to maintain oocyte quality during extended delays in fertilization, and while many components are identified, distinct phases have not been identified within oocyte RNP granules (Jud et al. 2008; Noble et al. 2008; Schisa, 2014).

In this study, we build upon our characterization of an imaging-induced stress that differentially modulates the condensation of RNA binding proteins in oocytes. While PGL-1 and the DEAD box helicase GLH-1 decondense in response to imaging stress, the KH domain protein MEX-3, DEAD box helicase CGH-1, and Lsm protein CAR-1 condense into granules (companion paper A). This differential stress response suggested that RNA binding proteins in large RNP granules of arrested oocytes may have distinct biophysical properties. We find that in large RNP granules, PGL-1 has liquid-like properties, and MEG-3 has gel-like properties, similar characteristics as in small P granules of early embryos. However, we find MEG-3 is not required for the condensation of PGL-1, CGH-1, or MEX-3 into large RNP granules which differs from its role in embryonic P granules. MEX-3 and CGH-1 may be distinct, intermediate phases of large RNP granules, as they remain stable at elevated temperature like MEG-3, but CGH-1 appears mobile via FRAP analyses, like PGL-1. We conclude the large RNP granules in arrested oocytes have distinct phases, and the phases correspond to the differential stress responses of RNA binding proteins and may have functional significance for gamete quality during extended meiotic arrest. Lastly, we identify PUF-5 as a potential regulator of condensation; however, it’s role in promoting the condensation of MEX-3 and MEG-3 into large RNP granules may be associated with the role of PUF-5 in oogenesis.

## METHODS

### Strains and maintenance

All worms were grown on nematode growth media using standard conditions at 20°C (Brenner 1974) unless specified. Strains used include: DG4269 mex-3(tn1753[gfp::3xflag::mex3]), JH3644 *fog-2(g71)* V; meg-3(ax4320)[meg-3::mCherry)]X, JH3269 pgl-1(ax3122[pgl-1::gfp]), JH1985 *unc-119(ed3); axIs1436[pCG33* pie-1prom:LAP::CGH-1], DG4215 puf-5(tn1726)[gfp::3xflag::puf-5]). Several strains were crossed into CB4108 *fog-2(q71)* as noted. Worms were synchronized using the hypochlorite bleach method.

### Microscopy and Image analysis

Worms were picked onto slides made with 2% agarose pads and paralyzed using 6.25mM levamisole. To image diakinesis oocytes of young hermaphrodites, synchronized 1 day post-L4-stage worms were used. To image meiotically arrested oocytes in *fog-2* strains, L4 females were separated from males and imaged 1 or 2dpL4 as indicated in figure legends.

Images were collected within 15 minutes of slide preparation unless otherwise noted. Imaging was done using a Nikon A1R laser scanning confocal microscope. All images for a given strain were collected using identical levels and settings. The number of granules in oocytes and total integrated density (intensity of fluorescence in granules) was determined using ImageJ particle analysis. To calculate the corrected total cell fluorescence (CTCF), ImageJ was used to measure the integrated density and area of the -1 to -5 oocytes, and the mean fluorescence of background. CTCF= integrated density – (area of selected ROI x mean fluorescence of background).

### Heat stress

The Tokai Hit stage top incubator was used at 34°C with the Nikon A1R laser scanning confocal microscope. Worms were placed on small NGM plates and incubated at 34°C in the stage top incubator for 40 minutes, or 2 hours as noted, before being transferred to a slide and imaged at 34°C within 15 minutes.

### Hexanediol assays

The hexanediol protocol of Updike et al., was adapted (2011). Worms were individually picked into 1 μL of egg buffer (25 mM Hepes, 120 mM NaCl_2_, 2 mM MgCl_2_, 2 mM CaCl_2_, and 48 mM KCl_2_) on a poly-L-lysine coated coverslip. The worm was dissected at both the proximal and distal end to release the gonad arms. Worms were imaged using the 60x objective every 2 seconds. After collecting baseline images (pre-treatment), 1ul of either egg buffer, 10% 1,6-hexanediol, or 1.0% SDS was added, and images were collected for 60 seconds (post-treatment). ImageJ was used to quantitate the number of granules and granule intensity in oocytes of pre- and post-treatment images. The ratio of pre/post values was graphed to determine if granules dissolved.

### Fluorescence Recovery After Photobleaching

For all strains, the 488 laser was used to photo-bleach an area of 3.68 µm^2^ within a single RNP granule in a -2 to -6 oocyte of a *fog-2* female. All proteins were imaged using a 500ms exposure and a 60x objective. Images were acquired every 8s during a recovery phase of 300s after photobleaching. The fluorescence intensities were normalized by dividing the intensity of the bleached zone by an unbleached fluorescent area (Hubstenberger et al. 2013). ImageJ was used to correct for movements during imaging. The Stacks-Shuffling-MultiStackReg plugin was used to align time series to an image at t=0.

### RNA-mediated interference

RNAi of *meg-3meg-4, mex-3*, and *puf-5* was performed by feeding using clones obtained from the Ahringer RNAi library or Openbiosystems library (Kamath & Ahringer, 2003). HT115 bacteria transformed with feeding vectors was grown at 37°C in LB + carbenicillin (conc.) for 5 hr, induced with 5 mM IPTG for 45 min, plated on RNAi plates [NGM (nematode growth media) + carbenicillin (conc) + IPTG (1mM)], and grown overnight at room temperature (Putnam et al 2019). Synchronized L1-stage worms were grown on NGM until the L4-stage when females were transferred to RNAi plates for ∼40 hr at 20°C. Controls for RNAi experiments included *unc-22(RNAi)* as a positive control for IPTG in RNAi media, and *lacZ(RNAi)* as a negative control since *lacZ* is not a *C. elegans* gene. All RNAi clones were sequenced to confirm the gene identities.

### Statistical analysis

Sample sizes were determined using G*Power 3.1 for power analyses; all experiments were blinded and done at least in triplicate. Data are presented as mean ± SEM unless otherwise indicated. Statistical analyses were performed on GraphPad Prism 9.1, and specific tests are noted in figure legends. P-values <0.05 were considered statistically significant.

### Data Availability

All strains are available at the *Caenorhabditis* Genetics Center (CGC) or on request from the Schisa lab or other *C. elegans* labs. The authors affirm that all data necessary to confirm the conclusions in this article are included in the article, figures, and tables.

## RESULTS AND DISCUSSION

### Differential responses to elevated temperature suggest distinct phases within large, oocyte RNP granules

In a companion paper (Elaswad et al., G3 submitted), we showed that imaging-induced stress triggers opposite phase transition-responses of PGL-1 and MEX-3/CGH-1/MEG-3. These results suggested the possibility that PGL-1 is a liquid phase of the large RNP granules in arrested oocytes, while MEX-3, CGH-1 and MEG-3 may be gel-like. Sensitivity to temperature changes can help distinguish liquid phases from solid phases (Weber and Brangwynne 2012; Banani et al. 2017); therefore, we exposed *fog-2(q71)* females with arrested oocytes to 34°C and monitored condensation in fluorescent reporter strains. In control *fog-2*; PGL-1::GFP females, large aggregates of PGL-1 granules were detected at the cortex among a heterogeneous mix of granules as expected (Jud et al. 2008) (Fig. 1A). In contrast, after 40 minutes at 34°C, while there was no statistical difference in overall numbers of granules, fewer large granules were apparent, and the total intensity of PGL-1 in individual granules was significantly reduced (Fig. 1B,C). In diakinesis oocytes of young hermaphrodites, PGL-1 also decondenses in response to heat stress (Jud et al. 2008; Fritsch et al. 2021; Watkins and Schisa 2021). Therefore, we conclude PGL-1 granules are sensitive to increased temperature in both diakinesis and arrested oocytes, consistent with a liquid phase. While these responses in oocytes are generally similar to the behavior of PGL-1 in early embryos, it is notable that PGL-1 disperses within 1 minute in a 2-cell embryo whereas significant dispersal was not detected in arrested oocytes until 40 minutes of temperature upshift. The germ granules in oocytes are composed of a different subset of proteins than embryonic germ granules, most granules are still attached to the oocyte nuclear envelope, and the viscosity of embryonic germ granules is much lower than that of RNP granules, all of which might contribute to the slower dynamics at elevated temperature (Updike and Strome, 2009; Brangwynne et al. 2009; Schisa, 2012; Hubstenberger et al. 2013). We next asked if MEG-3 is sensitive to elevated temperature and assayed *fog-2*; mCherry::MEG-3 females. We observed large granules in arrested oocytes of both control and 34°C worms (Fig. 1A), and no significant changes in granule number or intensity were detected after 2 hours at elevated temperature (Fig. 1B,C). Therefore, as in the embryo, MEG-3 seems stable at high temperatures in arrested oocytes, consistent with a gel-like phase.

**Figure 1.**
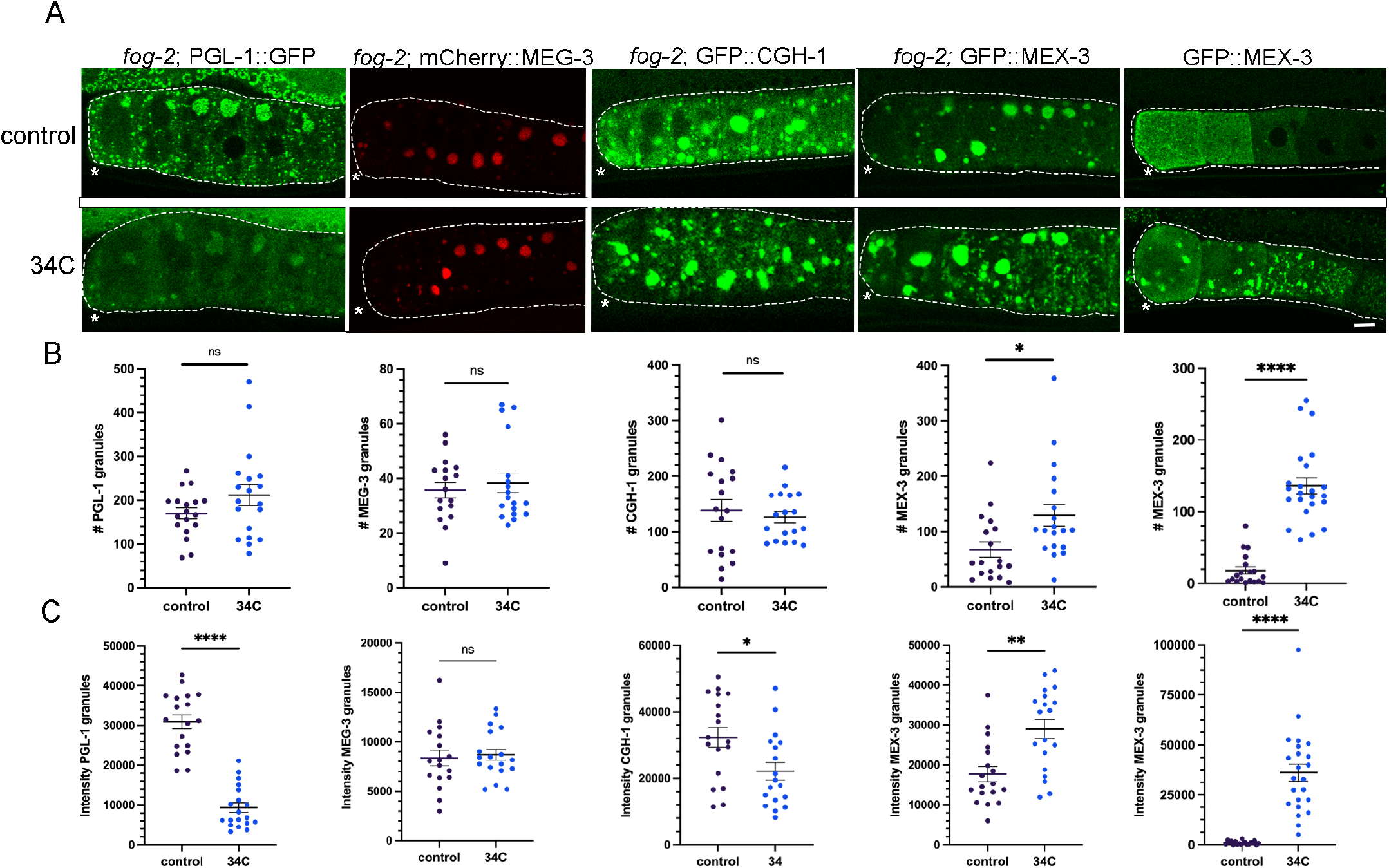
Increased temperature induces de-condensation and condensation of RNA binding proteins. (A) Micrographs of four fluorescently-tagged RNA binding proteins in a *fog-2* background, and GFP::MEX-3 in diakinesis oocytes. Top row: Arrested oocytes under control conditions of 20°C. Bottom row: Arrested oocytes after exposure to 34°C for 40 minutes (PGL-1) or 2 hours (all other proteins). Oocytes are outlined in white dashed lines. Asterisk marks the most proximal oocyte in each germ line. Scale bar is 10 µm. (B) Graphs showing the number of GFP granules in a single Z-slice of proximal oocytes (see Methods). For B and C, statistical significance was determined using the Mann-Whitney test. **** indicates p<0.0001, ** indicates p<0.01, * indicates p<0.05, ns indicates not significant. n=17-23. Error bars indicate mean ±SEM. (C) Graphs showing the total fluorescence intensity within granules in proximal oocytes (see Methods).

We next asked if CGH-1 is sensitive to elevated temperature by examining *fog-2*; GFP::CGH-1 females. We did not detect any changes qualitatively or find any significant changes in the number of granules; however, the intensity of GFP::CGH-1 in granules decreased slightly (Fig. 1A-C). Compared to PGL-1, CGH-1 appears only mildly sensitive to an upshift in temperature. Lastly, we asked if MEX-3 is sensitive to temperature changes. In control *fog-2*; GFP::MEX-3 females we detected large granules enriched at the cortex of arrested oocytes as expected (Jud et al. 2008) (Fig. 1A). After 2 hours at 34°C we observed a modest but significant increase in the number of granules, and significantly increased intensity of MEX-3 in granules (Fig. 1B,C). We therefore also asked if MEX-3 condenses in response to elevated temperature in diakinesis oocytes. In contrast to the modest change in MEX-3 condensation observed during imaging stress (Elaswad et al., G3 companion paper submitted), the majority of the MEX-3 protein that is normally dispersed throughout the oocyte cytosol appeared condensed into granules at the cortex or nuclear envelope (Fig. 1A,B). This result is consistent with a prior heat stress experiment performed using anti-MEX-3 antibodies on fixed worms (Jud et al. 2008). We conclude MEX-3 responds strongly to increased temperature by undergoing condensation in diakinesis and arrested oocytes. Taken together, the temperature shift experiments suggest PGL-1 may be the most liquid-like protein within large RNP granules, and this result correlates with PGL-1 being most sensitive to imaging-induced stress. In contrast, the condensation of MEX-3 and lack of sensitivity of MEG-3 to high temperature, suggest they may be gel-like phases in oocyte RNP granules.

### The PGL-1 phase within large oocyte RNP granules is liquid-like

To further characterize the phases of RNA binding proteins within large RNP granules, we next extruded gonads of *fog-2* females into the aliphatic alcohol hexanediol, a chemical shown to dissolve liquid phases (Patel et al. 2007; Lin et al. 2016). Hexanediol (HD) disrupts hydrophobic interactions, and induces the dispersal of the P granule proteins GLH-1 and PGL-1 in diakinesis oocytes (Updike et al. 2011). We analyzed the data by plotting the ratio of number of granules pre- and post-treatment, where no change would be indicated by 1.0. In egg buffer (EB)-treated control *fog-2;* PGL-1::GFP worms, we observed variable photobleaching in most worms, that contributed to a mean ratio of 0.54 (Fig. 2A,B). After exposing dissected gonads of *fog-2*; PGL-1::GFP to 5% HD, we observed a variable response and a decreased mean ratio of 0.36. In 55% of HD-treated germ lines, we observed a significant reduction in granule number (defined as a pre/post ratio ≤ 0.27), compared to only 11% of the control germ lines (Figure 2B). In some worms, we noted that the largest PGL-1 granules seemed less sensitive to HD treatment than smaller PGL-1 granules that more readily dissolved. Overall, the decrease in PGL-1 granule number in arrested oocytes suggests a similar sensitivity of PGL-1 within large RNP granules as for PGL-1 in the small P granules of diakinesis oocytes and embryos (Updike et al. 2011; Putnam et al. 2019).

**Figure 2.**
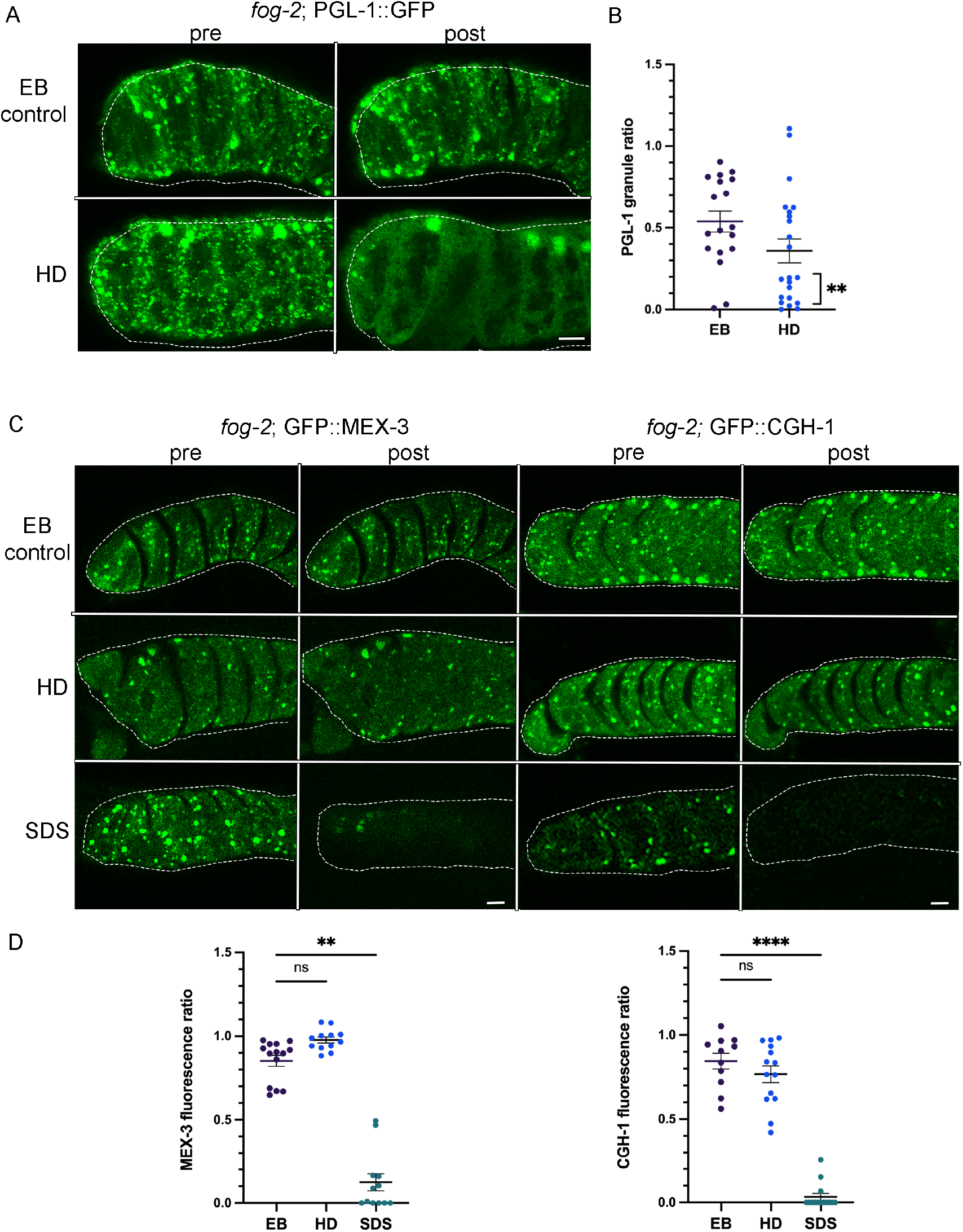
PGL-1 is a liquid-like phase within large RNP granules in arrested oocytes. (A) Micrographs of *fog-2*; PGL-1::GFP oocytes pre- and post-treatment with egg buffer (EB) control or 5% hexanediol (HD). Proximal oocytes are oriented to the left. Oocytes are outlined in white dashed lines. Scale bar is 10 µm. (B) Graph showing the ratio of number of PGL::GFP granules pre- and post-treatments. Statistical significance was determined using the Fisher Exact test. ** indicates p<0.01. n=18-22. (C) Micrographs of *fog-2*; GFP::MEX-3 and *fog-2*; GFP::CGH-1 oocytes pre- and post-treatment with egg buffer (EB) control, 5% hexanediol (HD), or 10% SDS. Scale bar is 10 µm. (D) Graphs showing the ratio of total intensity of GFP in granules, pre- and post-treatments. Statistical significance was determined using the Kruskal-Wallis test. **** indicates p<0.0001, ** indicates p<0.01, ns indicates not significant. n=11-14.

We next exposed dissected gonads of *fog-2*; GFP::MEX-3 and *fog-2*; GFP::CGH-1 worms to HD. In both strains, the granules appeared fairly stable with no significant change in response to HD as compared to the EB control (Fig. 2C,D). However, the granules did not appear to be irreversible aggregates because they rapidly dissolved upon exposure to the detergent SDS (Fig. 2C,D). One caveat to the interpretation of HD experiments is that they are not direct measures of liquid phases. Recent *in vitro* reports reveal that at 5-10% concentrations of HD, kinases and phosphatases can become inactive, and intermediate filaments dissolve, both of which could mask the properties of proteins (Lin et al. 2016; Duster et al. 2021). Nonetheless, these data are consistent with a liquid-like PGL-1 phase in RNP granules, and distinct phases of MEX-3 and CGH-1 that appear more gel-like.

### RNP granule proteins exhibit distinct dynamics in arrested oocytes

To determine RNA binding protein dynamics within RNP granules of arrested oocytes, we performed fluorescence recovery after photobleaching (FRAP) experiments. If a protein has liquid characteristics, it is expected to move randomly within an RNP granule. We determined the rates of exchange within a large granule by photobleaching a 3.68µm^2^ area in the center of large granules (Fig. 3A). We found that while PGL-1 and CGH-1 exhibited high rates of exchange, MEX-3 exhibited a somewhat lower rate of exchange (Fig. 3A, B). PGL-1, CGH-1, and MEX-3 all appeared to be mobile within RNP granules, similar to the complete exchange of the Lsm protein CAR-1 within RNP granules observed in prior FRAP experiments (Hubstenberger et al. 2013). Lastly, we tested MEG-3 and observed no intra-granule exchange of MEG-3 (the majority of trials) or extremely slow exchange dynamics (Fig. 3A, B). The rapid recovery of PGL-1 and lack of MEG-3 dynamics within granules were similar to the properties observed for granule-to-cytoplasm exchange in arrested oocytes (Putnam et al. 2019). The recovery rate we observed for PGL-1 within RNP granules was significantly faster than that detected for granule-to-cytoplasm exchange, consistent with the differential observed for CAR-1 (Hubstenberger et al. 2013), and suggesting that the interface between RNP granules and the cytosol is rate-limiting. Overall, our data build on prior results and establish distinct dynamics for four RNA binding proteins within RNP granules of arrested oocytes.

**Figure 3.**
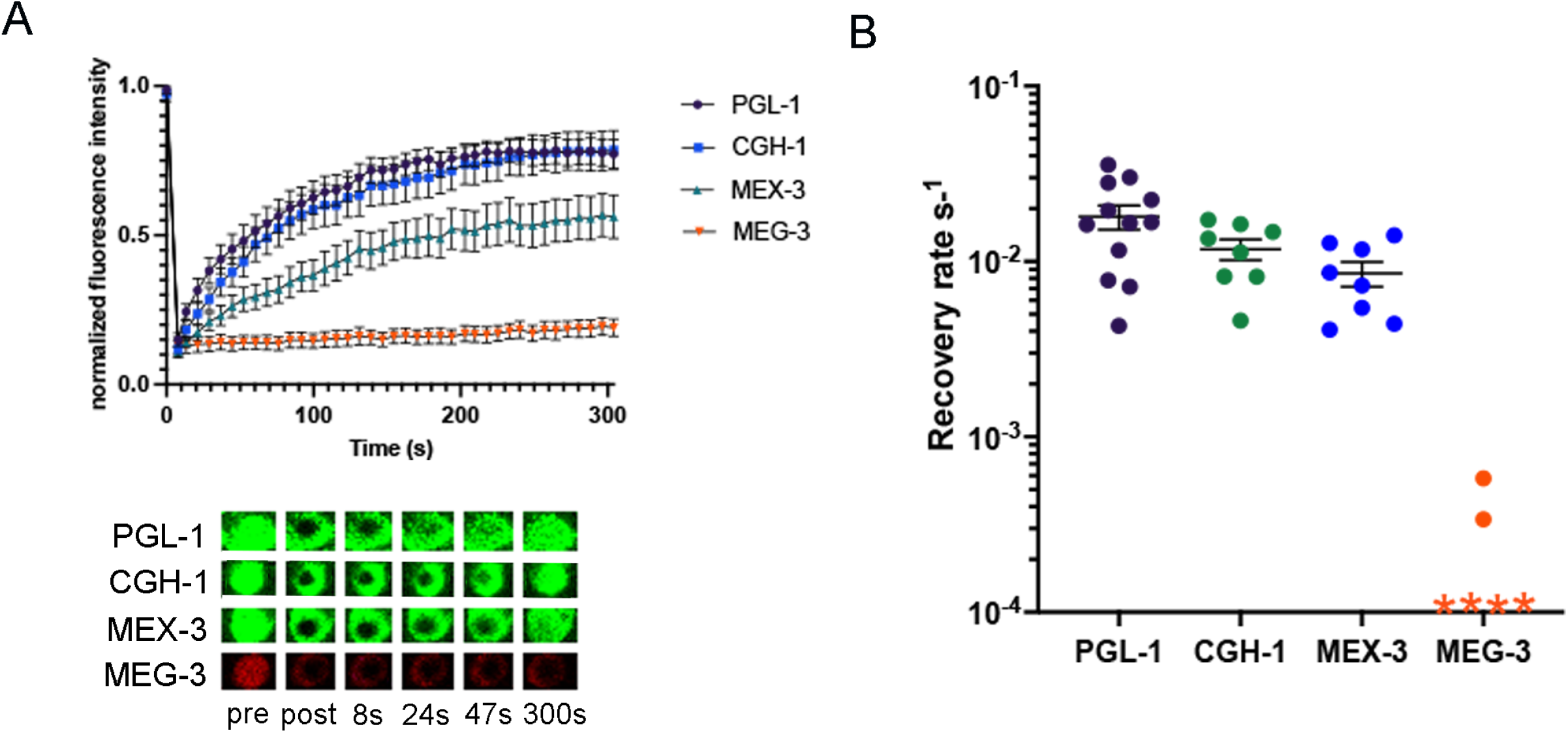
Oocyte RNP granule proteins exhibit distinct dynamics. (A) FRAP of PGL-1::GFP, GFP::CGH-1, GFP::MEX-3 and MEG-3::mCherry in arrested oocytes of *fog-2* females. An area of 3.68µm^2^ was bleached within a large granule. Granule intensity was measured every 7.8 seconds during a recovery phase of 300 seconds after photobleaching. Values were normalized to a reference granule intensity and plotted as an average of multiple experiments. (B) Graph shows recovery rates from FRAP experiments in A. Each symbol represents one FRAP experiment on one large RNP granule. Error bars are mean ±SEM. For MEG-3 data points indicated by an ‘*’, no recovery was detected. n=6-12.

### PUF-5 may promote condensation of MEX-3 and MEG-3 into large RNP granules of arrested oocytes

Given the stable and immobile behavior of MEG-3 within large, oocyte RNP granules, we hypothesized MEG-3 promotes the condensation of PGL-1, similar to its role as a stable scaffold of the liquid phases of germ granules in the early *C. elegans* embryo (Putnam et al. 2019). After RNAi of *meg-3* and its paralog *meg-4* in the *fog-2*; PGL-1::GFP strain, we observed large PGL-1 granules in arrested oocytes and high levels of fluorescence intensity in granules, with no detectable difference in PGL-1 condensation from the control RNAi worms (Fig. 4A). To ensure the *meg-3/meg-4* RNAi was efficiently decreasing expression of MEG-3, in parallel for each experiment we performed *meg-3/meg-4* RNAi in *fog-2*; MEG-3::mCherry females. We observed significant reduction of MEG-3 levels n all trials (Fig. 4A), demonstrating the depletion of *meg-3/-4* was effective. We next asked if MEG-3 is required for the condensation of CGH-1 or MEX-3, and observed no changes in the condensation of CGH-1 and MEX-3 into large granules after *meg-3/meg-4* RNAi compared to controls (Fig. 4A). We previously reported that MEG-3 appears to promote the condensation of MEX-3 into granules in arrested oocytes (Wood et al. 2016). The earlier set of experiments differed from the current experiments in two ways. First, the reporter strain for MEX-3 in the earlier experiments was made by microparticle bombardment, where the randomly integrated transgene is controlled by a *pie-1* promoter, and is non-rescuing (Jud et al. 2008); in contrast, here we are using an endogenously tagged allele of *mex-3* (Tsukamoto et al., 2017). Second, in the prior data set we reported a reduced amount of MEX-3 in granules after *meg-3* RNAi in 22% of worms, and the data were qualitatively analyzed (Wood et al. 2016). In contrast, here we used quantitative ImageJ-based particle analyses to more precisely assess the level of condensation of MEG-3. Taken together, we conclude that MEG-3 does not play a large role in promoting the condensation of PGL-1, CGH-1, or MEX-3 in meiotically arrested oocytes.

**Figure 4.**
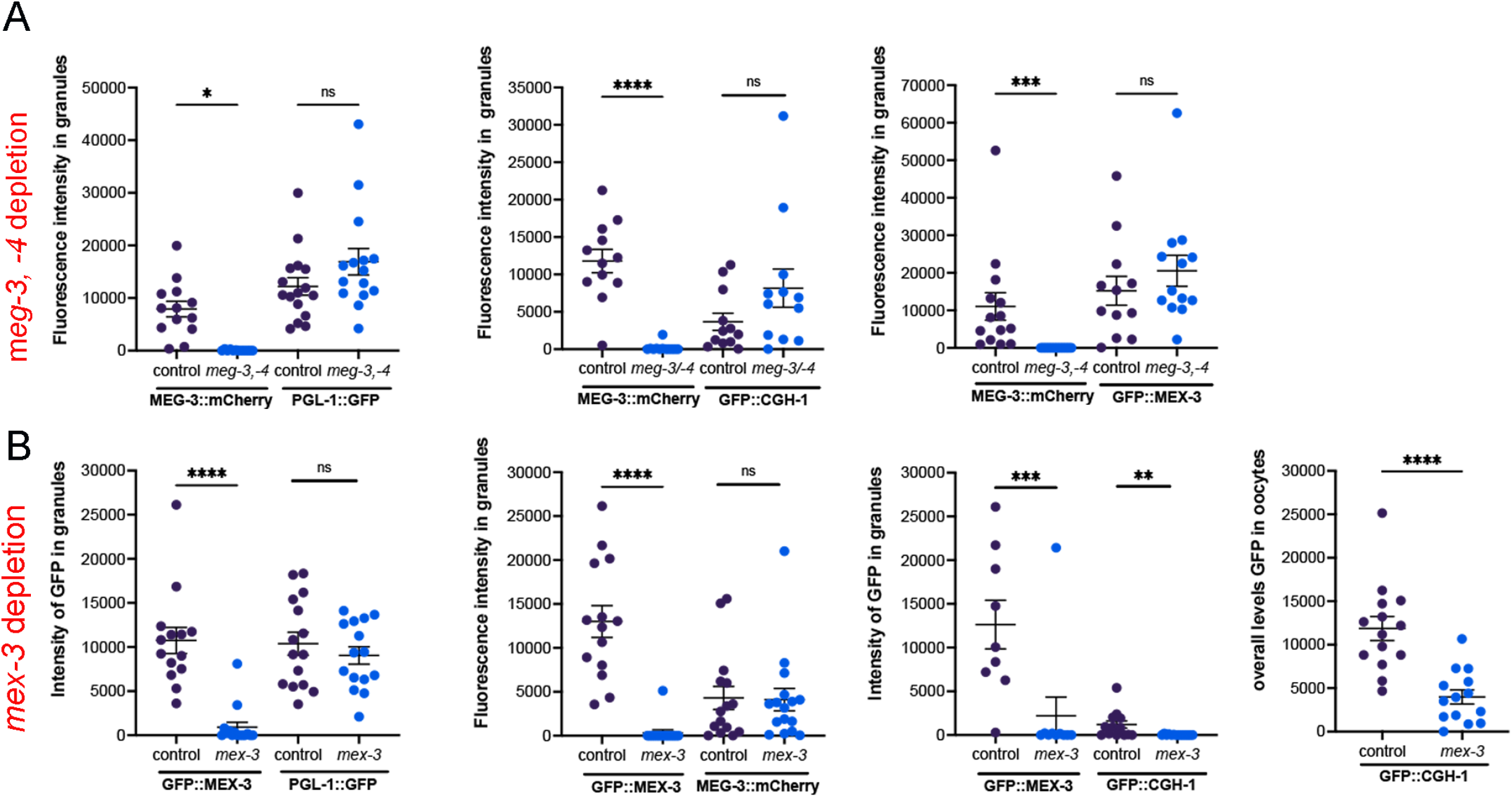
Neither MEG-3/MEG-4 nor MEX-3 are required to promote condensation of PGL-1 or each other into RNP granules. (A) RNAi of control *lacZ* or *meg-3 meg-4* was done by feeding in *fog-2* females with arrested oocytes. The MEG-3::mCherry strain was used as a positive control in each experiment to assess depletion of *meg-3*. The PGL-1::GFP, GFP::CGH-1 and GFP::MEX-3 reporters were used to assay condensation of the respective proteins. The intensity of fluorescence in granules is plotted on the Y axes. (B) RNAi of control *lacZ* or *mex-3* was done by feeding in *fog-2* females with arrested oocytes. The GFP::MEX-3 strain was a positive control in each experiment to assess depletion of *mex-3*. Reporter strains were used to assay condensation of PGL-1, MEG-3, and CGH-1. The intensity of fluorescence in granules, or the overall levels of fluorescence in oocytes (CTCF, see methods), is plotted on the Y axes. In all graphs, error bars are mean ±SEM. Statistical significance was determined using the Mann-Whitney test. **** indicates p<0.0001, *** indicates p<0.001, ** indicates p<0.01, * indicates p<0.05, ns indicates not significant. n=10-16.

Since MEX-3 condensed into granules in response to elevated temperature and was resistant to hexanediol treatment, we next hypothesized MEX-3 might be a gel-like phase that promotes condensation of PGL-1. After depletion of *mex-3* in the *fog-2*; PGL-1::GFP strain, we detected no significant difference in PGL-1 condensation compared to the control (Fig. 4B). The control *mex-3* RNAi in the *fog-2*; GFP::MEX-3 strain confirmed effective *mex-3* depletion in all trials. We next asked if MEX-3 promotes the condensation of CGH-1 or MEG-3 into large granules and detected no difference in condensation of MEG-3 (Fig. 4B). While we detected a significant reduction of CGH-1 condensed in granules, we also observed significantly decreased overall levels of CGH-1 in oocytes (Fig. 4B). These results suggest that MEX-3 may promote the translation or stability of CGH-1 protein in arrested oocytes. Consistent with the possibility of translational regulation, there are three MEX-3 recognition elements in the *cgh-1* 3’ UTR (Pagano et al. 2009). Overall, MEX-3 does not appear to have a major role in promoting the condensation of these three RNA binding proteins in arrested oocytes.

In a prior study, the PUF family of translational repressors was identified as a regulator of condensation of three RNA binding proteins in arrested oocytes. PUF-5 is reported to promote the condensation of two liquid-like RNA binding proteins, CAR-1 and PGL-1, into large granules (Hubstenberger et al. 2013); and depletion of *puf-5/-6/-7* prevents condensation of CGH-1 in fixed samples. Therefore, we asked if PUF-5 promotes the condensation of MEX-3 in arrested oocytes. In all trials, RNAi-mediated depletion of *puf-5* was very effective in the control GFP::PUF-5 strain (Fig. 5A,B). After RNAi of *puf-5* in *fog-2*; GFP::MEX-3 worms, we observed two distinct phenotypes. In ∼40% of *puf-5* RNAi oocytes, we detected large MEX-3 granules similar to the *lacZ* RNAi control result; however, in ∼60% of oocytes MEX-3 appeared at higher levels throughout the cytosol than in the control and only small granules were detected (Fig. 5A,B). In 8 of 15 gonads the fluorescence intensity in granules was below the lowest level seen in the control (Fig. 5B). This result suggests PUF-5 may play a role in promoting MEX-3 condensation; however, a caveat is a correlation between reduced condensation and increased disorganization of the germ line. The PUF-5/-6/-7 proteins have redundant roles in oocyte differentiation. Depletion of *puf-5,-6,-7* results in abnormally small oocytes that are often stacked into two or three rows; however, depletion of *puf-5* alone results in normally differentiated and organized oocytes (Lundlin et al., 2007). In the *fog-2* background, RNAi of *puf-5* alone appears to result in a partially penetrant, disorganization phenotype where oocytes are stacked in overlapping rows instead of a single, linear row (Fig. 5A, transmitted light), and the decondensation of MEX-3 could be due to the disorganization and therefore an indirect effect of *puf-5*.

**Figure 5.**
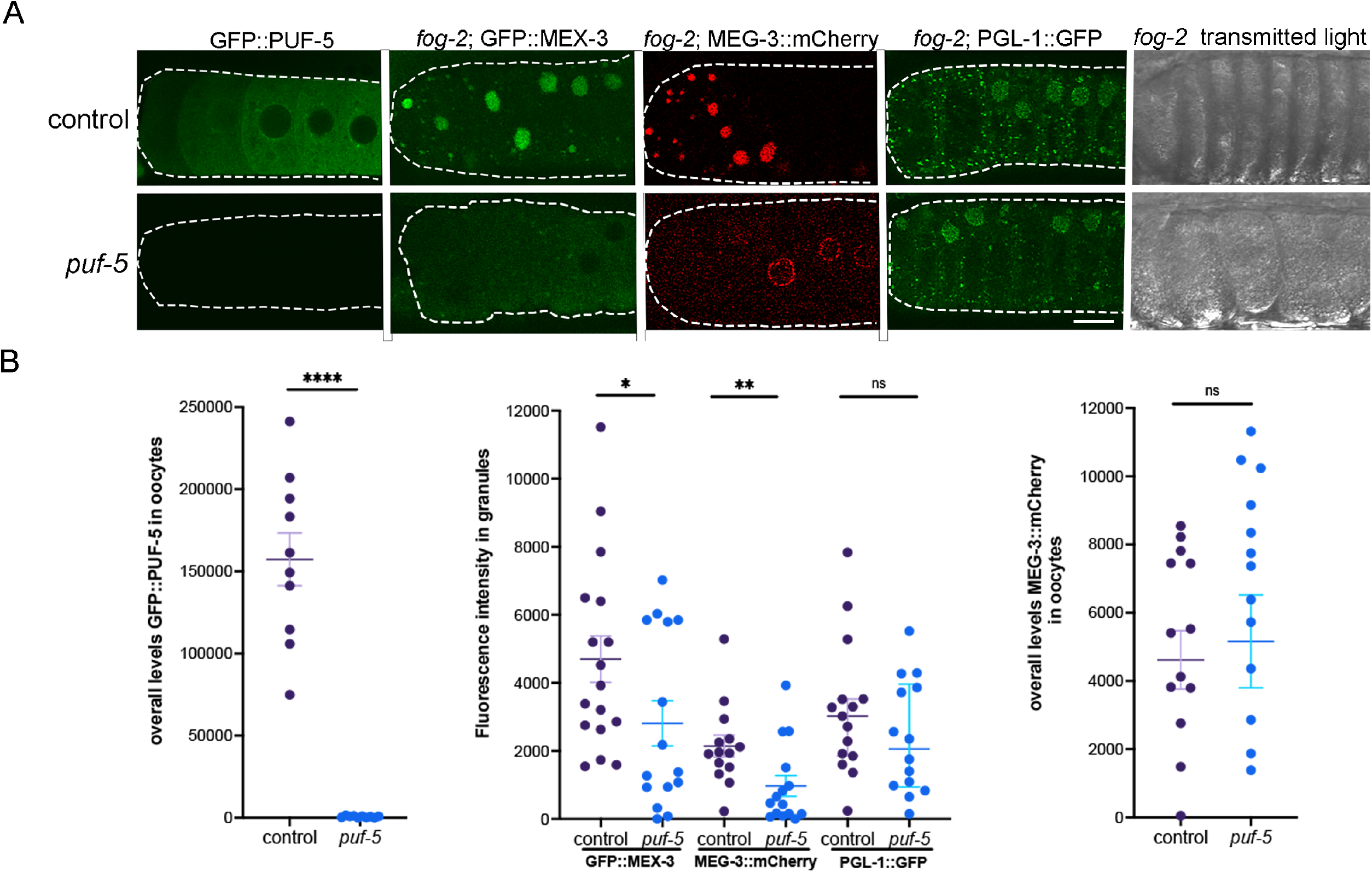
PUF-5 may promote condensation of MEX-3 and MEG-3 into large granules. (A) Micrographs of GFP::PUF-5 in diakinesis oocytes, three fluorescently-tagged RNA binding proteins in a *fog-2* background, and *fog-2* oocytes visualized by transmitted light. Depletion of control *lacZ* or *puf-5* was performed by feeding RNAi. The GFP::PUF-5 strain was used as a positive control in each experiment to assess the extent of depletion of *puf-5*. The GFP::MEX-3, MEG-3::mCherry, and PGL-1::GFP reporters were used to assay condensation of the respective proteins in arrested oocytes. Oocytes are outlined in white dashed lines, with the most proximal oocyte on the left. Scale bar is 10 µm. (B) Graphs showing the overall levels of GFP::PUF-5 or MEG-3::mCherry in oocytes (CTCF, see methods), or the intensity of fluorescence in granules, as plotted on the Y axes. Statistical significance was determined using the Mann-Whitney test. **** indicates p<0.0001, ** indicates p<0.01, * indicates p<0.05, ns indicates not significant. n=10-22. Error bars indicate mean ±SEM.

We next asked if PUF-5 regulates MEG-3 condensation in arrested oocytes. In the negative control, the majority of MEG-3 protein was detected in large, bright, cortical granules as expected (Fig. 5A). After *puf-5* depletion in ∼40% of gonads, large, cortical MEG-3 granules were detected similar to the control, but in ∼60% of gonads MEG-3 was detected at increased levels diffuse throughout the cytosol and in small, perinuclear granules (Fig. 5A). The condensation of MEG-3 into large granules was significantly decreased after *puf-5* depletion, and there was no significant difference in overall levels of MEG-3 in oocytes (Fig. 5B). Therefore, the data suggest PUF-5 promotes the condensation of MEG-3 into large, cortical granules in arrested oocytes. However, as for MEX-3, this role may be indirect as it is associated with the partially penetrant disorganization phenotype (Fig. 5A).

Lastly, we assayed PGL-1 condensation using a strain where the endogenous *pgl-1* locus is tagged with GFP (Putnam et al., 2019), expecting to reproduce the result of decondensed PGL-1 after depletion of *puf-5* (Hubstenberger et al., 2013). In the control, we observed a mix of small and large PGL-1::GFP granules in arrested oocytes as expected. After *puf-5* depletion, the large majority of worms appeared similar to control worms, and no significant difference in the amount of PGL-1 condensation was detected (Fig. 5 A,B). Although negative results in RNAi experiments must be interpreted cautiously, *puf-5* RNAi appeared to be very effective in depleting *puf-5* expression in the control GFP::PUF-5 worms (Fig. 5A); therefore, we conclude that PUF-5 does not play a major role in promoting condensation of PGL-1. We cannot be sure why our result differs from the previously published result; however, the earlier experiment used a strain with a randomly integrated transgene made by microparticle bombardment in which the *nmy-2* promoter drives expression of a PGL-1::mRFP translational reporter, RNAi was done by microinjection rather than feeding, and no quantitative analyses were described to measure variability or oocyte disorganization after *puf-5* RNAi (Hubstenberger et al. 2013; Wolke et al. 2007). Overall, we conclude PUF-5 may selectively regulate the condensation of RNA binding proteins into RNP granules in arrested oocytes; however, the decondensation phenotypes may be indirectly caused by germ line disorganization or partially penetrant resumption of meiotic maturation.

## Conclusions

Taken together, our results here demonstrate that large RNP granules of arrested oocytes have distinct phases (Table 1; Fig. 6). In prior studies, cytological results have suggested substructure within the large RNP granules of arrested oocytes. For example, MEX-3 and PAB-1 (poly(A) binding protein) appear to occupy sub-domains of granules in fixed samples, as do PGL-1 and *pos-1* RNA, and live imaging of reporter strains shows PGL-1 and CAR-1 occupy subdomains of RNP granules (Schisa et al. 2001; Jud et al. 2008; Hubstenberger et al. 2013). Our current results build on the cytological observations of substructure in oocyte RNP granules with multiple, functional studies that indicate PGL-1 is dynamic and a liquid-like phase within oocyte RNP granules (Table 1; Fig. 6). The liquid behavior of PGL-1 correlates with the sensitivity and de-condensation of PGL-1 in response to imaging-induced stress (Table 1). The FRAP data for CGH-1 showed strongly dynamic behavior within large RNP granules. These data were somewhat surprising since large granules of CGH-1 in arrested oocytes were not sensitive to imaging stress, nor strongly sensitive to elevated temperature or hexanediol treatment (Table 1; Elaswad et al., G3 submitted). CGH-1 may be an intermediate phase of large RNP granules, distinct and less liquid-like than PGL-1. Our results also demonstrate that MEX-3 and MEG-3 are less dynamic phases of oocyte RNP granules (Fig. 6). MEG-3 has the slowest dynamics via FRAP, and MEX-3 demonstrates the strongest condensation phenotypes at elevated temperature. Both proteins may be gel-like phases of RNP granules; however, neither appears to strongly promote the condensation of other RNA binding proteins in RNP granules. In contrast, we identified PUF-5 as a potential regulator of MEX-3 and MEG-3 condensation into large granules in arrested oocytes, adding to its previously characterized roles in promoting condensation of CAR-1, and CGH-1 (Hubstenberger et al. 2013). Future studies should resolve if the observed *puf-5* condensation phenotypes are caused by a resumption of meiotic maturation or germ line disorganization. Given the role of RNA in recruiting RNA binding proteins to stress granules (Campos-Melo et al. 2021), investigations of the role of RNA as possible nucleators or molecular scaffolds of oocyte RNP granules will also be important to gain a more complete understanding of RNP granule formation.

**TABLE 1.** Differential protein dynamics in large RNP granules suggest distinct phases.

|  | Oogenesis stage | PGL-1 | CGH-1 | MEX-3 | MEG-3 |
| --- | --- | --- | --- | --- | --- |
| Imaging stress* | diakinesis | decondense | condense | condense | not tested |
|  | arrested | decondense | stable | stable | stable |
| Temp. shift | arrested | decondense | partially decondense | condense | stable |
| Hexanediol | arrested | dissolve | stable | stable | not tested |
| SDS | arrested | not tested | dissolve | dissolve | not tested |
| Mobility (FRAP) | arrested | mobile | mobile | moderately mobile | less mobile |
| Dynamics |  | ++++ | +++ | + | - |
| Phases |  | Liquid-like | Mix of liquid-like and gel-like | Mostly gel-like | Gel-like |

\*Companion paper, Elaswad et al., G3 submitted

**Figure 6.**
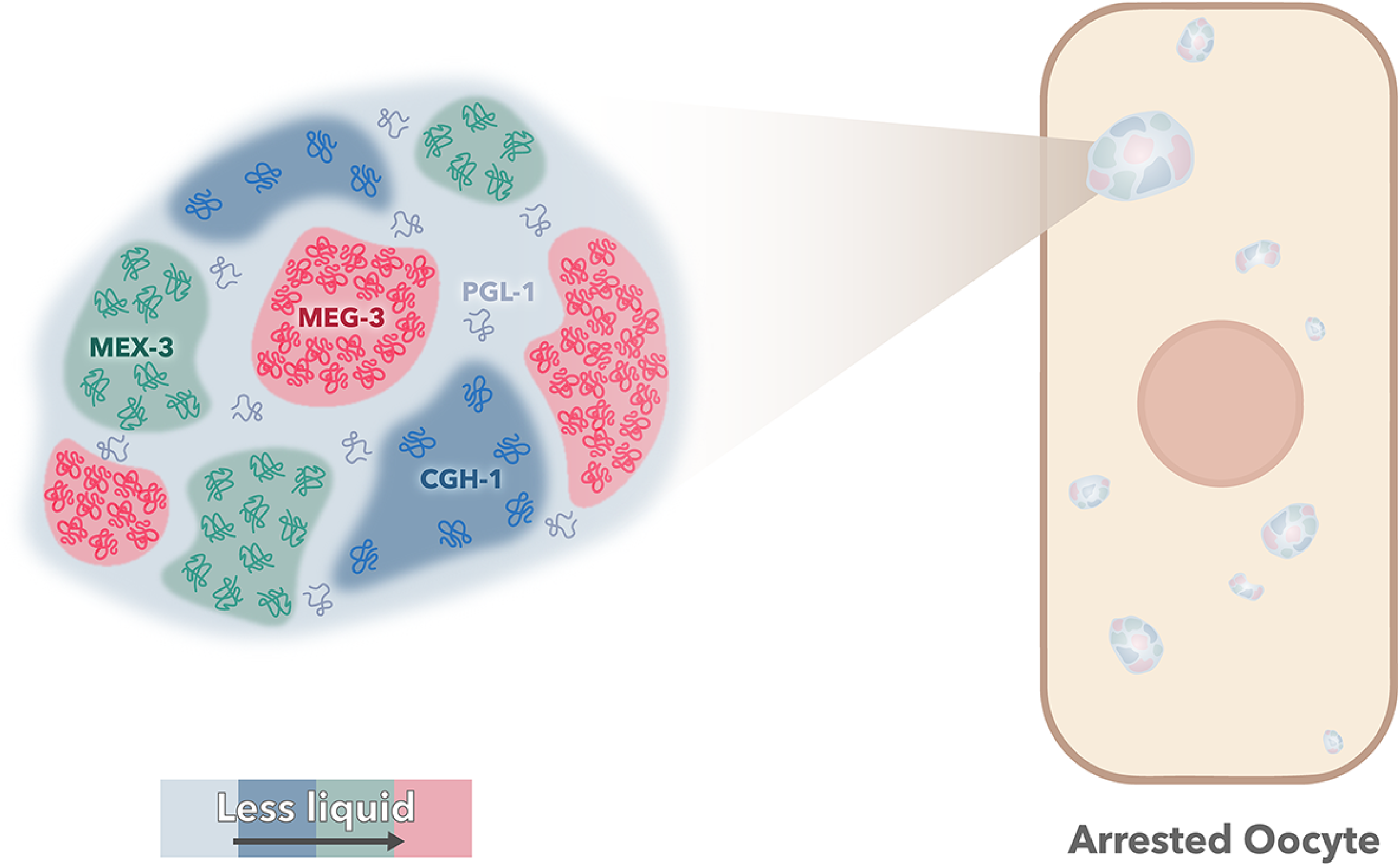
Model of distinct phases of RNA binding proteins in RNP granules. The relative dynamics of the four RNA binding protein phases within large RNP granules is indicated by color. PGL-1 is the most liquid-like (light blue), CGH-1 has a mix of liquid and gel-like properties (dark blue), MEX-3 has gel-like properties (green), and MEG-3 appears the most gel-like (red).

Our data demonstrate the properties and regulators of RNA binding proteins within large, oocyte RNP granules have some similarities, but are not identical, to small germ granules of early embryos. The differences may be influenced by the significant differences in granule size, composition, viscosity, or differential function. Ultimately, gaining a better understanding of the phases of RNP granules in arrested oocytes should lead to insights into the function of the condensed RNA binding proteins when fertilization is delayed an extended time, and oocyte quality needs to be maintained.

## ACKNOWLEDGEMENTS

We appreciate strains provided by Dr. Dustin Updike, Dr. Geraldine Seydoux, and the *Caenorhabditis* Genetics Center, which is funded by NIH Office of Research Infrastructure Programs (P40 OD010440). We thank Dr. Updike for sharing protocols and helpful discussion. We thank Wormbase. C.P. was supported by an Undergraduate Summer Scholars grant from the Office of Research and Graduate Studies at Central Michigan University. Funding for this work came from a CMU Faculty Research and Creative Endeavors grant to J.S. and NIH grant 2R15GM109337-02A1 to J.S.

